# Directed differentiation of hPSCs through lateral plate mesoderm for generation of articular cartilage progenitors

**DOI:** 10.1101/2021.03.24.436807

**Authors:** Christopher A Smith, Paul A Humphreys, Mark A Naven, Fabrizio E Mancini, Susan J Kimber

## Abstract

Developmentally the articular joints are derived from lateral plate (LP) mesoderm. However, no study has produced LP derived prechondrocytes or preosteoblasts from human pluripotent stem cells (hPSC) in a chemically defined manner. Differentiation of hPSCs through the authentic route, via an LP-osteochondral progenitor (OCP), may aid understanding of human cartilage development and the generation of effective cell therapies for osteoarthritis. We refined our existing chondrogenic protocol, incorporating knowledge from development and other studies to produce a LP-OCP from which prechondrocytes- and preosteoblast-like cells can be produced. Results show the formation of an OCP, which can be further driven to prechondrocytes and preosteoblasts. Prechondrocytes cultured in pellets produced cartilage like matrix with lacunae and superficial flattened cells expressing lubricin. Additionally, preosteoblasts were able to generate a mineralised structure. This protocol can therefore be used to further investigate cartilage development and in the development of joint cartilage for potential treatments.

## Introduction

Osteoarthritis (OA) is a painful degenerative disease affecting millions worldwide [1]. It is caused by the continued destruction of the articular cartilage, a tissue lining the ends of joints which enables their smooth movement, however; the underlying bone is also affected. This degeneration results in inflammation which further exacerbates the disease. Ultimately the effect of OA is pain for the sufferer, a reduction in mobility and an overall decrease in their quality of life, as well as great cost to society [2, 3]. Articular cartilage has a low regenerative capacity and although artificial prostheses may be offered, a tissue engineered replacement of the damaged tissue involving cellular therapies is an attractive option. Human pluripotent stem cells (PSC) offer a great opportunity for the generation of cellular therapies for degenerative diseases. Their proliferative potential and ability to differentiate into any cell in the body [4] including chondrocytes for cartilage treatment, offers them distinct advantages over other cell types such as mesenchymal stromal cells (MSCs), which, whilst able to produce chondrogenic cell types [5], are limited in expansion and therefore bulk supply of cells [6] and can have donor-specific issues with heterogeneity, quality and efficacy [7].

Importantly, in the majority of OA cases, both the articular cartilage and the underlying bone (forming the osteochondral system) are affected. Osteochondral constructs are therefore required to aid in the healing of such defects, whilst they are likely to improve the integration of cartilage [8]. Consequently, a cell population able to produce both cartilage and bone of the articular joint or “articular progenitor” in an experimentally controlled manner, would be particularly useful for joint TE, to minimise protocol complexity and maximise healing potential.

Our group have developed directed differentiation protocols to produce chondroprogenitors from hPSCs [9, 10], with updated protocols replacing BMP4 with BMP2 to improve chondrogenesis [11], or substituting small molecules for growth factors to remove batch variability [12]. However, though these differentiation pathways can produce chondroprogenitors, they are not able to produce osteogenic cells, and only give moderate quality of cartilage pellets when cultured further (unpublished). One way of ensuring the production of cells present during the formation of the fetal tissue is to attempt to recapitulate the development process more closely [13]. The differentiation of hPSCs to chondrocytes has, to date, relied heavily on differentiation through the paraxial mesoderm pathway, generating chondrocytes able to produce recognizable cartilage like matrix [14–17]. Whilst chondrocytes can be derived through the paraxial mesoderm/ somite pathway, these cells do not typically populate the limb bud to generate the joints and their articular cartilage. As such, utilising this route may result in a different chondrocyte phenotype, for example expressing type-X collagen and high levels of RUNX2 indicative of mineralizing prehypertrophic non-hyaline cartilage [15]. Indeed some hPSC differentiation protocols instead rely on minimal use of differentiating reagents and isolating the appropriately differentiated cells instead of directing a more homogenous differentiating population [18, 19] or through the isolation of cells from mixed aggregates such as from embryoid bodies [20–22]. However, the cells which produce the cartilage and bone comprising the joints, originate from the limb-bud, which develops from lateral plate mesodermal tissues [23]. Unlike somite derived cartilage in the body, this developmental pathway gives rise to the unique niche from which the articular cartilage arises [24]. As such, this differentiation pathway should give rise to a stable hyaline cartilage phenotype, and likely generates chondrocytes with differences to those produced by the paraxial route [25] which gives the axial skeleton in development. Though there have been attempts to generate chondrocytes from cells deriving from lateral plate [20, 26] they use undefined methods, such as formation of embryoid bodies rather than utilise directed differentiation [13]. More recently Loh *et al* have detailed the generation of lateral plate mesoderm derivatives from hPSCs, with indications of limb bud tissue [27]. Building on our previous protocols and this additional knowledge, our study aims to recapitulate the developmental system of limb-bud formation to produce an osteochondral progenitor, which can give rise to both prechondrocytes and preosteoblasts in a directed manner. To this end we define the serum free **R**efined **A**rticular **P**rogenitor **I**ntegrated **D**ifferentiation (RAPID) protocol.

## Materials and Methods

### hPSC Cell culture

For continued pluripotent culture, hPSC lines Man-13 [28] and Man-7 [11] were cultured in feeder-free culture conditions in 6-well plates coated with vitronectin (5 ug/mL) (Thermo) in mTSeR1 (Stem Cell). Cells were passaged at 70-80% confluence with 0.5 mM EDTA supplemented with revitacell (Gibco). Medium was initially changed after 24 hours to remove revitacell, then subsequently every 2 days.

### RAPID progenitor differentiation protocol

hPSCs were split onto fibronectin coated (16.6 ug/mL) (Millipore) 6-well plates at 100,000 cells/cm^2^ and cultured till 60-70% confluence before the start of each protocol. Upon reaching confluence, pluripotency medium was removed, and directed differentiation basal medium (DDBM)(See supplementary data) added subject to particular differentiation pathway as described in full in Figure 1A. Briefly, for osteochondral lineages, cells were directed using our Refined Articular Progenitor Integrated Differentiation (RAPID) protocol, at first culturing cells in CHIR99021 (3mM), Activin A (1ng/mL), BMP2 (as BMP concentration is crucial in lateral plate development, and to avoid batch variation, each BMP batch activity was titrated and used at ECx66), FGF2 (20ng/mL) and PI3K inhibitor; PIK90 (100nM) to generate a culture expressing mid primitive streak markers (days 1-2) [27, 29]. At day 3 only the growth factors BMP2 (10ng/mL) and FGF2 (10ng/mL) were continued but with the addition of WNT inhibitor C59 (100nM) and ALK5 inhibitor SB431542 (2mM), to drive cells towards lateral plate mesoderm (days 3-4 orange lineage Fig 1A). At days 5-6 cultures were subjected to a CHIR pulse to induce limb-bud-like mesoderm or withheld to promote cardiac-like mesoderm (blue lineage Fig1A). Limb-bud mesoderm progenitors were further induced either through application of DDBM medium containing GDF5 (40ng/mL) and BMP2 (0.5xECx66) to prechondrocyte cells (day 7-11, green lineage Fig1A), or using osteogenic-basal medium (OM) (See supplementary data) containing CHIR (2mM) to preosteoblast cells (days 7-8, red lineage Fig1A) (7-8). Separately, day 5 lateral plate mesoderm progenitors were further differentiated to cardiac mesoderm and into cardiomyocytes through days 8-28 using Basal-cardio Medium (BM) (Lateral plate derived [LPD] cardiac protocol, blue lineage Fig1A). For the full application sequence of growth factors and small molecule see Figure 1B. For media composition see supplementary data.

**Figure 1.**
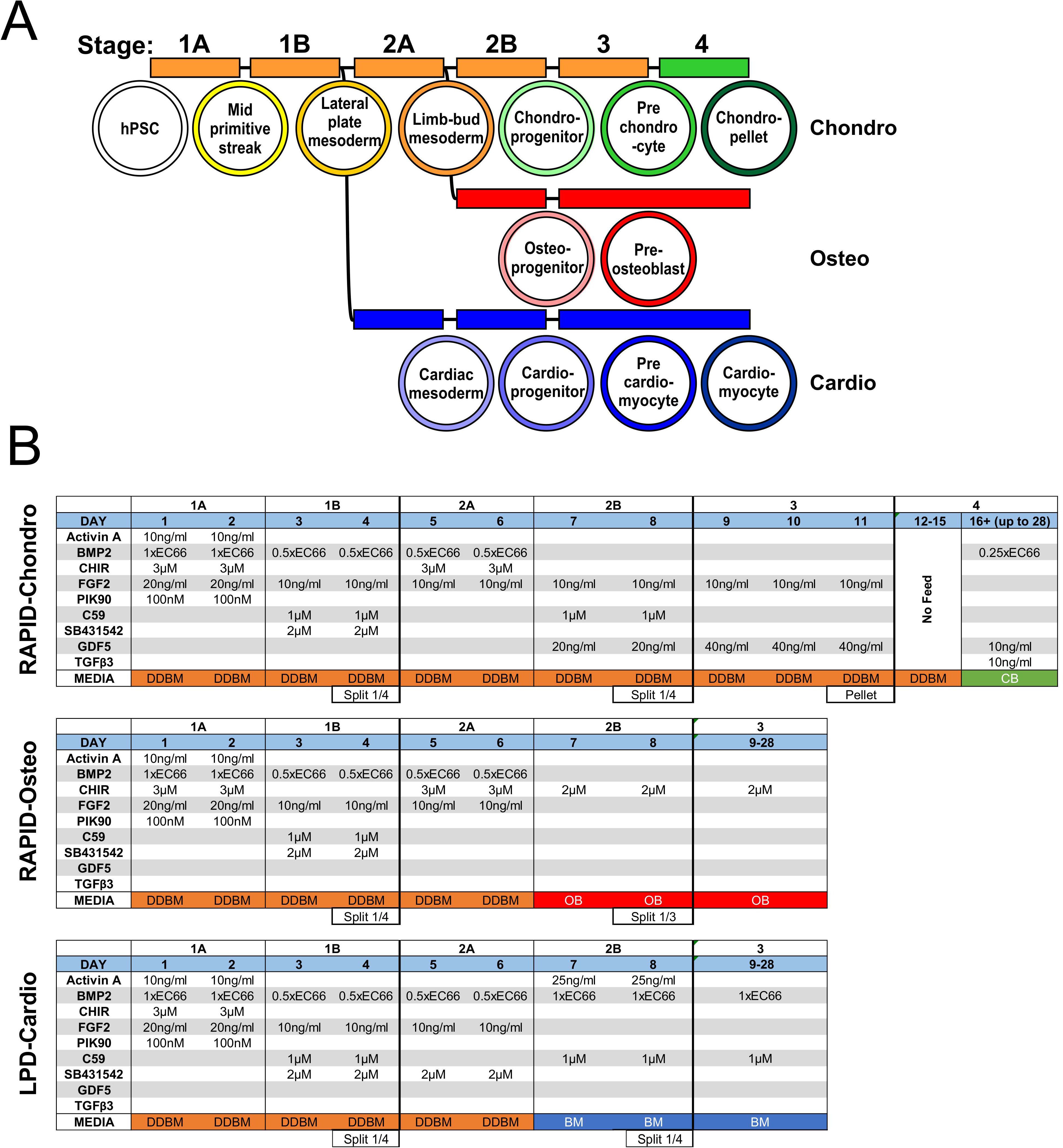
RAPID protocol. A) Flow diagram of stages and media required to achieve all 3 cell types during the RAPID (inc LPD). B) Growth factor regimen for RAPID (inc LPD) protocols at each day for the chondrogenic, osteogenic and cardiogenic protocols.

### Chondrogenic pellet culture

Day 11 chondroprogenitors were removed from tissue culture plastic and centrifuged at 300 xRCF in day 11 medium for 5 minutes to sediment cells. Caps were left loose to enable gaseous transfer and tubes incubated for 3 days at 37°C to facilitate the production of a spherical pellet, as detailed previously [12]. At day 11+3, medium was changed for chondrogenic-basal medium (CM) containing GDF5 (10 ng/mL), TGFβ3 (10 ng/mL) and BMP2 (0.5x ECx66). For full composition of media see supplementary data. Pellets were cultured for up to 28 days with medium changed every 3-4 days.

### Osteogenic culture

Day 8 osteoprogenitors were removed from fibronectin coated plastic and split 1:3 into non-coated 6-well plates. Cells were cultured with OM supplemented with 2μM CHIR changed every 3-4 days up to day 28. Additional samples were continued to day 28 for matrix staining.

### QRT-PCR Gene transcriptional analysis

Samples were taken from both 2D and 3D culture as described previously [12]. Samples were transferred to RLT buffer directly and RNA extracted by use of a RNeasy QiAgen kit. Extracted RNA was converted to cDNA using the ABI-RT kit (Life technologies). Quantitative real-time polymerase chain reaction (qRT-PCR) was conducted using PowerUp™ SYBR™ green (Life technologies) and assayed for genes associated with stages of development and eventual cell types (Chondrogenic, osteogenic, cardiogenic) (see supplementary data for full primer sequences). Data was analysed and displayed as expression relative to housekeeping gene *GAPDH*.

### Alizarin red staining

Monolayer samples were washed in PBS, fixed in 4% PFA solution for 10 minutes at room temperature, then washed 3 times in deionized water. A 1% (W/V) Alizarin solution was added to the wells and incubated at room temperature for 10 minutes, then washed in deionized water until all excess dye had been removed. Samples were then air dried before images were taken under bright field using a vertically mounted camera (Nikon D330).

### BCIP/NBT active ALP enzymatic stain

Samples were fixed in 4% PFA at room temperature for 2 minutes, then washed 3 times in PBS. Sufficient BCIP-NBT was added to cover the cell monolayer and incubated at 37°C for 20 minutes. The solution was removed then the monolayer was washed with deionized water 3 times and allowed to dry. Images were taken under bright field using a vertically mounted camera (as above).

### Histology

Chondrogenic pellets were fixed in 4% PFA overnight at 4°C then transferred to 70% ethanol before processing and embedding in PFFE blocks. Embedded samples were sectioned at 5μm thickness. Sections were stained for H&E, Picrosirius Red or Alcian Blue.

For immunohistochemistry, sections were subject to 0.1M citrate antigen retrieval at 95°C for 10 mins before the primary antibody for type-II collagen (Millipore, MAB8887), ACAN [30] or lubricin (Abcam, ab28484) was added. Samples were incubated overnight at 4°C, then treated with either fluorescent secondary antibodies (Thermo) and ProLong mountant (Thermo) for visualisation under a microscope (Olympus IX71), or with a biotin conjugated secondary antibody (R&D) and visualised using 3,3′-Diaminobenzidine (DAB) and images taken on a slide scanner (3D Histech Panoramic250).

### Statistical analysis

All statistical analysis was run using Prism Graph-pad. Samples were tested for normality using a D’Agostino-Pearson test. Statistical significance was calculated using Kruskal-Wallis, Mann-Whitney or Wilcoxen (indicated in figure legend). A p-value of ≤ 0.05 was considered as statistically significant.

## Results

We set out to generate chondroprogenitors which would develop further to form joint-like cartilage though the lateral plate mesoderm route. We named the developed protocol Refined Articular Progenitor Integrated Differentiation (RAPID) protocol and checked its authenticity by inducing other lateral plate derived differentiation derivatives from early-stage cells.

### The generation of osteochondral progenitors through lateral plate mesoderm

In the first step ActivinA, FGF2, BMP2 and the GSK3 inhibitor CHIR 99021 (to replace Wnt3a) were used to generate mid-primitive streak like cells as employed previously by ourselves [12] and others [27]. Additionally, the PI3K inhibitor PIK90 was used to help inhibit endodermal differentiation [31]. QRT-PCR gene transcriptional analysis indicated a clear increase in the primitive streak associated genes Brachyury (T) [32] and MIXL1 [33], and mid/posterior primitive streak CDX2 was present at day 2 of the differentiation protocol (Figure 2A) [34]. Following this and the removal of ActivinA and CHIR 99021, CDX2 expression was present during cell maturation towards mesodermal tissue together with high HAND1 and HAND2 expression indicating lateral plate specification [27]. Additionally, transient TBX6 transcripts were observed during the differentiation to mesodermal tissue[35] TBX6 subsequently diminished at day 6 as expected. Importantly the transcription factors FOXF1, NKX2.5 and ISL1 were also significantly upregulated by day 4. These, together with CDX2, are essential for lateral plate mesoderm specification [34, 36]. During the mesodermal and subsequent stages there was also no significant increase in the paraxial mesoderm associated marker MEOX1, and PAX1 was significantly decreased by day 6, indicating significant paraxial mesoderm derived populations were unlikely to be differentiating (Fig 2C).

**Figure 2.**
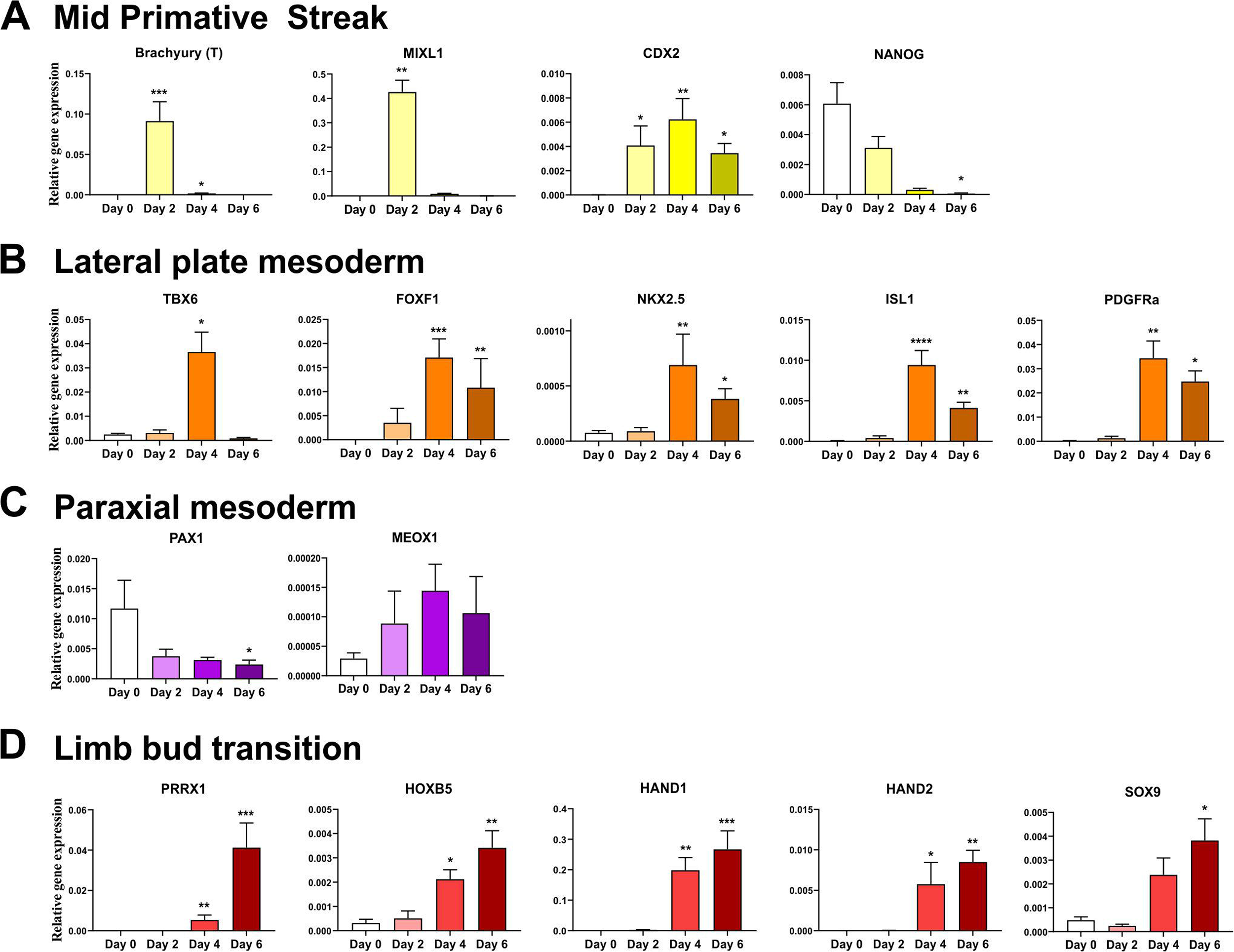
hPSC are directed through lateral plate lineage to limb-bud like progenitor cells. Gene expression relative to GAPDH using 2-delta ct +SEM, for transcriptional analysis of hPSCs differentiating into mid-primitive streak (Day2) (A), lateral plate mesoderm (Day4) (B), paraxial markers (C), and limb-bud transition like cells (Day 6) (D). D’Agostino-Pearson test was used to test for normality. All series tested using Kruskal-Wallis test to determine statistical significance. Significance relative to hPSC (day 0) control cells (+ ≤0.05, ++≤0.01, +++≤0.001, ++++≤0.0001). (N=6 biological repeat)

Following a second application of CHIR 99021 (day 5-6) on day 6, there was a decrease in the expression of lateral plate mesoderm markers (Figure 2B), which coincided with an increase in limb-bud specification markers PRRX1, HOXB5, HAND1 and HAND2 as well as SOX9 (Figure 2D). Additionally, the pluripotency/primitive steak marker NANOG was significantly decreased by this stage (Fig 2A). These day 6 cells expressed PRRX1, PDGFRa and SOX9 indicating specification to an osteochondral progenitor (OCP) [23, 24], and were then subsequently used for further chondrogenic or osteogenic differentiation.

### Transition from limb-bud like osteochondral progenitors to prechondrocytes

The lateral plate derived limb-bud like cells were driven in their differentiation to chondrogenic progenitors with inhibition of the Wnt pathway with C59 and the addition of the articular joint growth factor GDF5. At day 8 chondroprogenitor cells expressed significant levels of the transcription factors SOX5, SOX9 and ARID5B as well as the ECM components ACAN, COL2A1 and COL1A2 compared to day 0 (Figure 3A). Cells continued in the development with additional GDF5 till day 11 and resulted in sustained expression of the chondrogenic markers, however PRG4 (lubricin) was not expressed at either day. A further hPSC line, Man7, was also subjected to chondrogenic differentiation using the RAPID protocol, and also displayed significant expression of the chondrogenic markers SOX5, SOX9, ARID5b, COL2A1, and ACAN as well as COL1A2 at day 11 (Supplementary data Figure S1). These prechondrocytes were then taken into pellet culture to test their chondrogenic differentiation ability.

**Figure 3.**
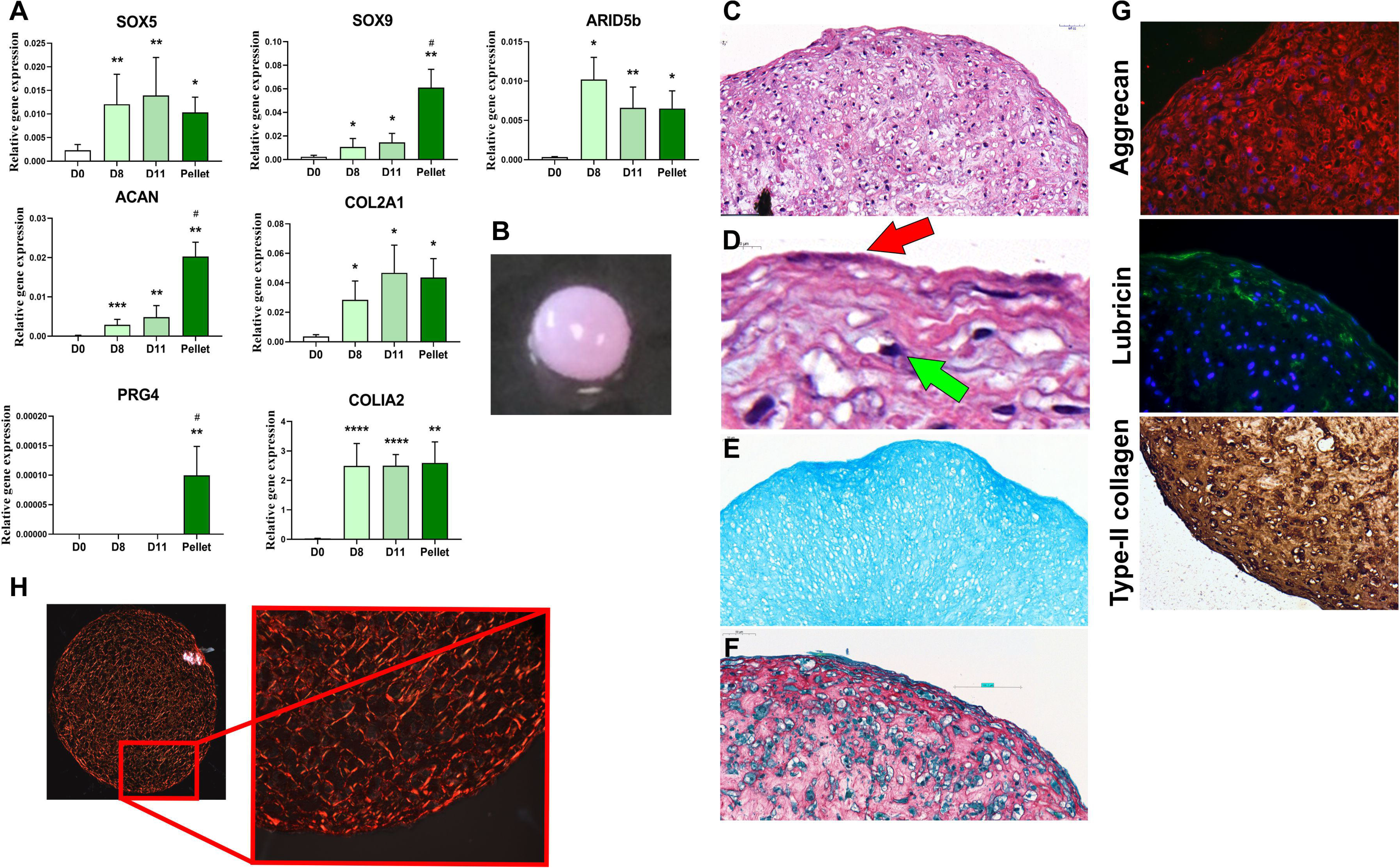
Differentiation of chondroprogenitors from limb-bud like progenitor cells. A) QRT-PCR analysis of chondrogenic gene expression during differentiation in 2D (until day 11) and in 3D pellet culture (day 11+14). Gene expression data displayed as relative to housekeeping gene GAPDH +SEM. Mann-Whitney test was used to determine statistical significance relative to hPSCs (day 0) control cells (+≤0.05, ++≤0.01, +++≤0.001). # indicates significant difference to day 11 (N=5 biological repeat). B) Image at day 11(2D) +28 d in 3D for pellets and enlarged insert. C) H&E histological stain of day 11+28 pellet, and D) enlarged view of H&E including arrows for flattened surface cells (red arrow) and cells in lacune (green arrow). E) Alcian blue stain of day11+28 pellet. F) picrosirius red collagen matrix stain. G) Immunofluorescence images for day 11+28 pellets stained for Aggrecan (red), Lubricin (green) and type-II collagen (IHC). In all images DAPI was used for visualisation of cell nuclei. H) Polarised light imaging of picrosirius red staining of day 11+28 pellet.

### Prechondrocytes can form cartilage like matrix

To test the chondrogenic potential of the prechondrocytes, day 11 cells were pelleted and cultured for up to 28 days. This further differentiation in 3D pellet culture resulted in the continued high-level expression of SOX5, ARID5B, COL2A1 and COL1A2, with significant increases in SOX9 and ACAN compared to day 11. Importantly, in 3D pellet culture cells also expressed the articular surface proteoglycan PRG4 (lubricin). Type-X collagen was not detected during 2D or 3D pellet culture (data not shown), indicating the cells were not developing into pre-hypertrophic chondrocytes.

After 28 days in 3D pellet culture, pellets displayed a spherical, translucent, glossy appearance similar to the articular cartilage surface (Figure 3B). Histological staining displayed a cartilage-like structure (Figure 3C) with cells present in lacunae (Figure 3D, green arrow on insert), and surface cells showing a flattened elongated morphology (Figure 3D, red arrow). The pellets stained for Alcian blue, with a stronger intensity towards the surface (Figure 3E), whilst Picrosirius red staining indicated a dense collagen matrix (Figure 3F). Indeed, polarised light imaging after Picrosirius red staining indicated fibrillar collagen networks throughout the pellets, with a change in orientation towards the surface of the pellets (Figure 3H), reminiscent of the expected change in cell alignment at the cartilage surface. Histological analysis of these pellets identified the key cartilage components aggrecan, lubricin and type-II collagen within the structure (Figure 3G). Whilst aggrecan was present throughout the pellet structure, lubricin was only present near the surface of the pellet. Additionally, type-II collagen staining was strong throughout the structure of the pellet, with darker staining towards the pellets surface.

Whilst cells from day 8 cells of the differentiation protocol can be used to produce pellets, in our experience those pellets are fragile, tend to disintegrate after a few days and are noticeably less stable than those produced from cells at day 11. Therefore, day 11 cells were used for the formation of pellets.

### Transition from limb-bud-like osteochondral progenitors to preosteoblasts

To assess the osteogenic potential of the limb-bud-like progenitor cells, differentiation was continued after day 6 of the above protocol in osteogenic medium. Gene expression analysis showed significant increases in expression of the osteogenic transcription factors SPARC (osteonectin) and RUNX2 by day 8 in comparison to day 0 (Figure 4A). This significant increase was sustained at day 14. The essential bone ECM component type-I collagen was also significantly increased by day 8 and remained high at day 14, with significant expression of the osteoblast-expressed ECM component, BGLAP (osteocalcin), by day 14. No type-X collagen was detected at any stage (data not shown). Differentiating preosteoblast cells were cultured till day 28 and showed continued significant expression of osteoblast markers compared to day 0. Additionally, their matrix mineralisation ability was assessed by staining of the resulting structure with Alizarin red, which resulted in an orange/red colouring of the produced matrix (Figure 4B) including nodes apparent in the matrix (insert). This indicated the formation of inorganic calcium rich deposits. Additionally, BCIP-NBT staining produced a strong blue colour throughout the matrix, suggesting active ALP enzyme activity in the matrix/cells.

**Figure 4.**
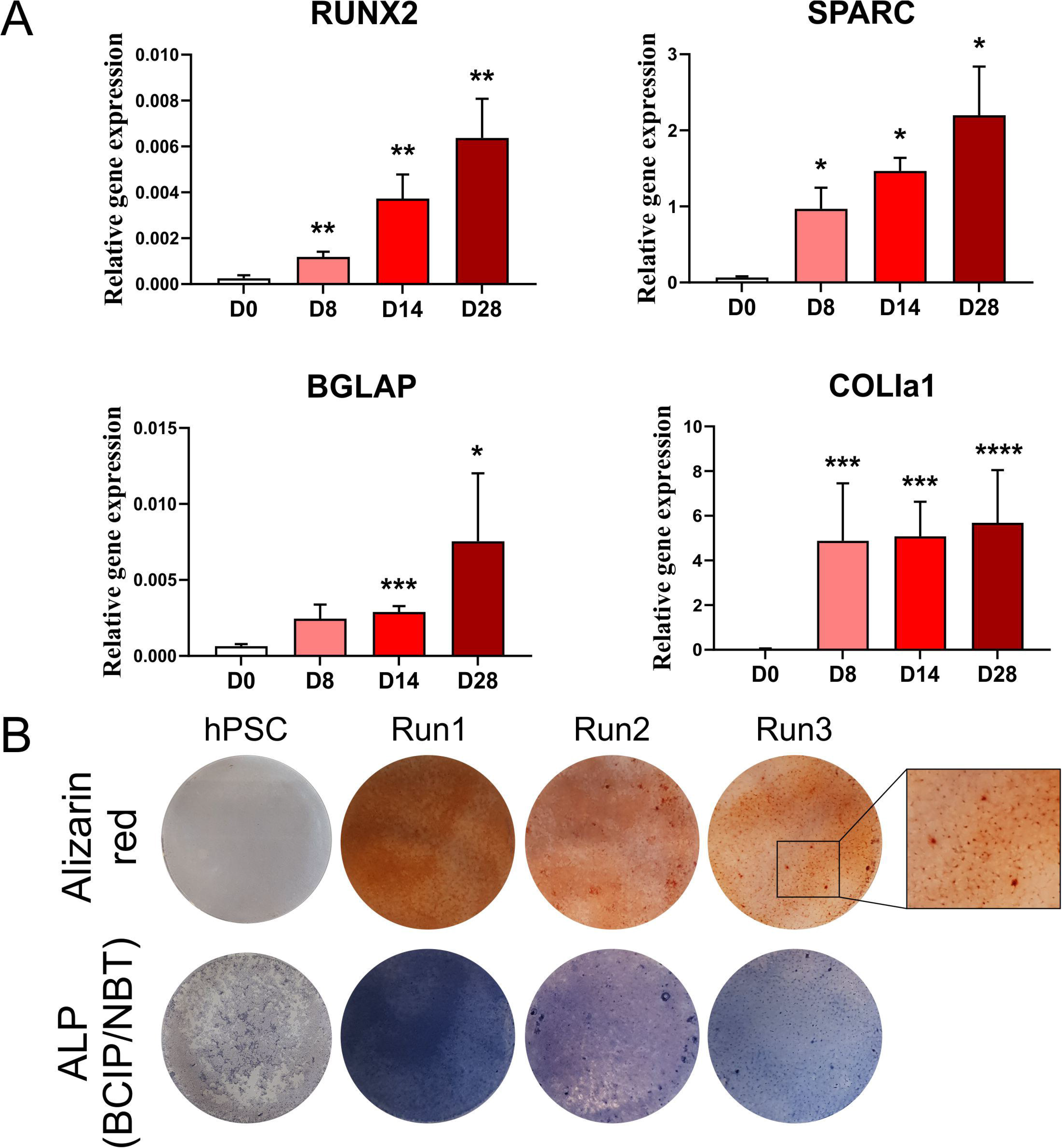
Differentiation of preosteoblast like cells from limb-bud-like progenitor cells. A) QRT-PCR gene expression analysis during osteogenic differentiation pathway up to day 28. Gene expression data displayed as relative to housekeeping gene *GAPDH* with SEM. Mann-Whitney test was used to determine statistical significance. Significance relative to hPSC (day 0) control cells (+ ≤0.05, ++≤0.01, +++≤0.001) (N=5 biological repeat day 0 to 14, N=3 biological repeat for day 28). B) Alizarin red mineralisation assay for preosteoblast like cells cultured till day 28 (N=3 biological repeat). C) BCIP-NBT active ALP enzyme assay for preosteoblast like cells cultured till day 28 (N=3 biological repeat).

### Cardiomyocyte derivation from lateral plate mesodermal cells

To confirm our route to chondrocytes and osteoblasts we tested if cardiomyocyte progenitors could be generated from day 4 lateral plate-like cells (Figure 1) as cardiomyocytes also originate from the lateral plate mesoderm. Instead of promoting a limb-bud like cell, day 4 lateral plate-like cells were cultured on, without a CHIR pulse at day 5-6, to generate a cardiac-like mesoderm. The resulting day 6 cells were then transferred to a cardiac mesoderm differentiation cocktail in BM for up to 28 days (Figure 1A). Gene transcription analysis of day 6 samples showed a significant increase in the cardiac marker NKX2.5, the cardiac specific TNNT2 transcript, as well as cardiac maturation transcription factors GATA4 and 6 [37] (Figure 5). These cardiac markers remained significant up to day 28 (Figure 5). Cells formed clusters around day 11, which began beating asynchronously (Supplementary video). By day 28 these clusters had increased in size and were more synchronous in their beating.

**Figure 5.**
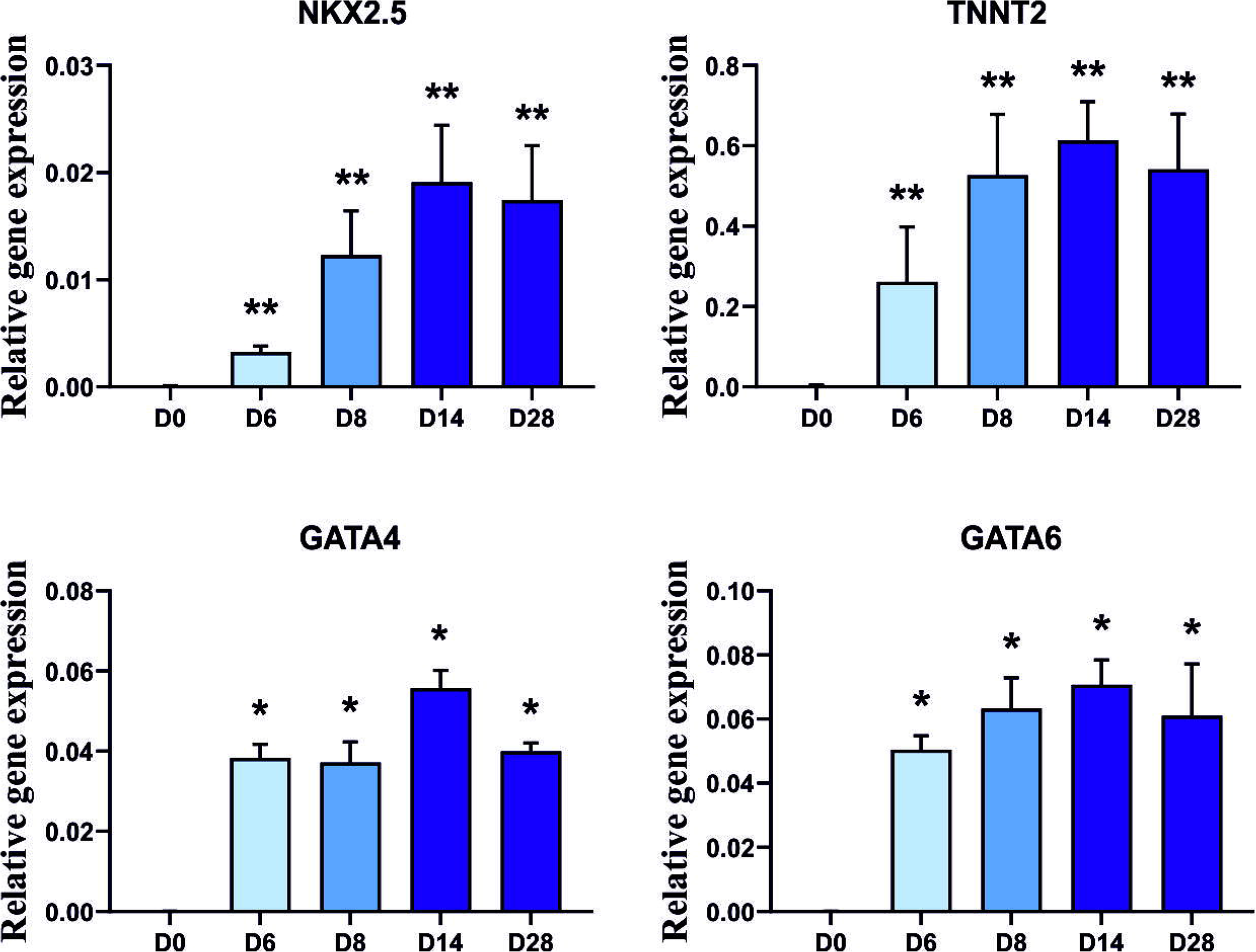
Differentiation of cardiomyocyte progenitor cells from lateral plate lineage through cardiac mesoderm. Gene transcriptional analysis of cardiomyocyte genes during differentiation to beating cardiomyocytes. (For video of structures contracting please see supplementary evidence). Mann-Whitney test was used to determine statistical significance. Significance relative to hPSC (day 0) control cells (+≤0.05, ++≤0.01) (N=5 biological repeat for days 0 to 14, N=4 for day 28).

### RAPID protocol shows improved chondrogenic gene expression to standard DDP

We tested the chondrogenic potential of the lateral plate derived RAPID prechondrocytes compared to those of the original DDP [7]. Paired RAPID and DDP samples were differentiated and their expression of the key chondrogenic factors were compared at the end of the respective 2D protocols. Day 11 RAPID prechondrocytes showed increased gene expression compared to Day 14 DDP cells, with significant increases in transcription factors SOX5 and SOX9 as well as the matrix components COL2A1 and ACAN (Figure 6A). Additionally, we also compared the histology of pellets produced from the DDP [12] with those of the RAPID prechondrocytes. RAPID prechondrocytes derived pellets are considerably larger than those from DDP and had a deeper alcian blue staining (Figure 6B). Whilst both pellet types stained for the main matrix components, the RAPID pellets displayed greater staining for type-II collagen and aggrecan, whilst maintaining a stronger type-II collagen stain towards the pellet surface.

**Figure 6.**
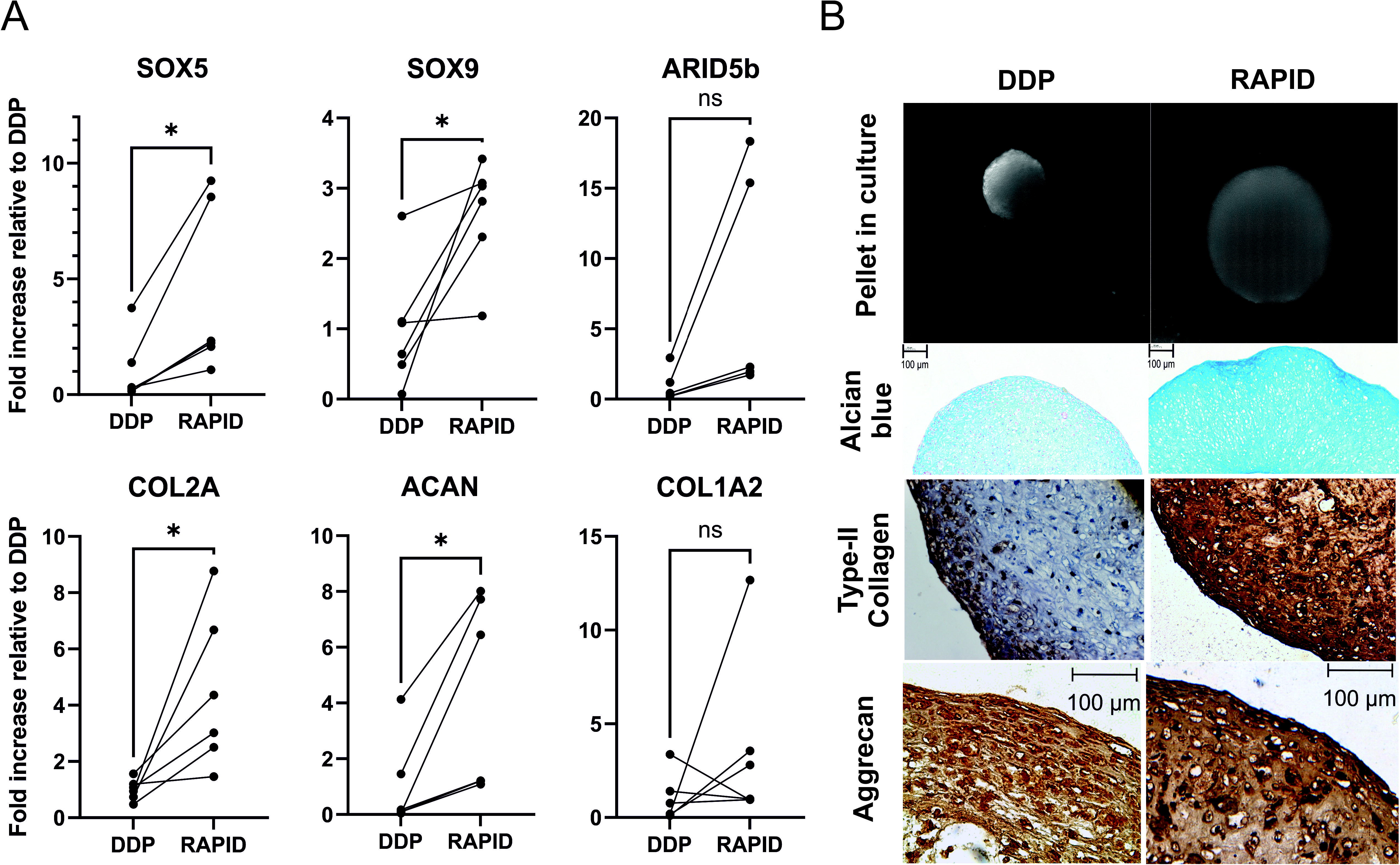
Comparison in chondrogenic gene expression between paired existing DDP and lateral plate derived RAPID cells. Man7 PSCs were differentiated to chondroprogenitors through both the DDP and RAPID protocols. A) QRT-PCR gene expression assessment for the main transcription factors and ECM components in pair match DDP and RAPID samples. DDP samples were taken at day 14 compared to day 11 in the RAPID protocol. Data is expressed as normalised to existing DDP protocol, with bars indicating paired experiments. B) Light microscopy image of pellets, alcian blue matrix staining, type-II collagen and aggrecan staining in 2D+28 days pellets. A Wilcoxen test was used to determine statistical significance. + indicate significance to hPSC (day 0) control cells (+ ≤0.05, ++≤0.01) (N=6 biological repeat).

## Discussion

In this study we have detailed the use of an improved, defined protocol (RAPID), to generate the lateral plate derived cell types prechondrocytes, preosteoblasts and cardiomyocyte cells from hPSC. This protocol relies on knowledge of development to achieve this; differentiating the cells first through mid-primitive streak, then lateral plate mesoderm from which limb bud OCPs can be derived, and further differentiated into both prechondrocytes and preosteoblasts. Furthermore, beating cardiomyocyte-like cells can be derived from a generated cardiac mesoderm intermediate.

Though other groups have reported the production of lateral plate mesoderm from hPSCs [27], their focus has been on specifying cardiac progenitors and they did not differentiate limb-bud derived tissues such as osteochondral progenitors. Other studies, which aimed at generation of articular chondrocytes, have achieved chondrogenesis through the use of EB differentiation [21, 22, 26]. However, in this study we have modified our previous defined differentiation protocol (DDP) [9, 11, 12] influenced by the invaluable early lineage data of Loh *et al* [27], and here detail the growth factors and small molecule requirements for production of these cells types from the Man l3 and 7 lines. The time-specific growth factor applications were influenced by our previous differentiation protocol, and those of others [27, 34], including adjacent mesoderm lineages such as kidney [29], which we have previously utilised to generate kidney organoids [38]. Similar to our original differentiation pathway, these protocols all indicate the importance of early application of ActivinA, FGF and CHIR as surrogate for Wnt stimulation. However, we utilised BMP2 in our protocol, having previously identified it as able to induce hPSC-chondrogenic gene expression [11], maintaining this throughout the protocol to form our OCPs alongside a CHIR pulse at days 5-6. Importantly, we have also included the activity of BMP2 required in our differentiation protocols. The derivation of lateral plate requires a careful balance of BMP signalling. As such, this growth factor activity data is aimed at making our protocols more reproducible to reduce the issue of batch/ supplier variation in BMP2 activity.

Using this protocol we can produce articular OCPs in as little as 6 days, with cells expressing PRRX1, SOX9 and PDGFRa as found in the developing limb [23, 24]. Continuing their differentiation from OCP to chondroprogenitor and ultimately a prechondrocytes by day 11, we again followed in the natural articular joint development. Crucially, to achieve this we utilised GDF5, a critical signalling molecule in the natural development of articular cartilage [39–42], and which we have used previously [11, 12, 43]. Importantly, we also maintained the cells in GDF5 between day 8 and 11, which was required to ensure a robust stable structure, similar to our previous findings [11].

When limb-bud/ articular progenitor cells were differentiated to prechondrocytes, there were significant increases in expression of chondrogenic transcription factors and ECM genes. Crucially, pelleting led to the formation of smooth translucent tissue, which displayed cells in a lacuna structure. Whilst we have previously detailed the production of chondrogenic progenitors and cartilaginous pellets from prechondrocytes [12], this current protocol is faster and generates pellets more similar to hyaline cartilage by following a lateral plate pathway. In paired samples, the RAPID produced chondroprogenitors expressed significant increases in the transcription factors SOX5 and SOX9, as well as the matrix components COL2A1 and ACAN. Importantly, the pellets produced from the RAPID protocol are larger, with more uniform structure and much stronger Alcian blue, GAG, staining. Crucially to articular chondrogenesis, our RAPID protocol produced pellets stained for lubricin on the outer region of the pellets, which together with a more flattened cell phenotype, a change in matrix orientation, and an increased fibrillar collagen, indicates a similarity to the superficial zone in articular cartilage [44, 45]. These observations indicate the production of cartilage with distinct zones, a feature that will be crucial for cell engineering in replacement of damaged cartilage [44, 46].

Importantly, OCPs did not express significant PAX1 and MEOX1 indicating they were not paraxial and not developing through a somite lineage. Though differentiation through paraxial mesoderm and somites has frequently been used to produce chondrocytes [13] ultimately appropriate directed chondrogenic differentiation through the limb-bud-like mesoderm is likely to produce cells more akin to the native articular chondrocytes generated during development from growth plate progenitors and interzone cells [39]. This route may affect the cells’ maturation and long-term phenotype, as articular cartilage is not produced through the somite chondrogenic development pathway. The cells produced through the RAPID protocol were positive for the major chondrogenic factors as well as the surface zone proteoglycan lubricin (PRG4) [46, 47], indicative of permanent, hyaline cartilage. This is in comparison to type-X collagen, which was not evident in our system but which is expressed in mineralising cartilage such as the interface with bone, the growth plate and endochondral ossification during bone healing.

In addition to prechondrocytes, the OCPs differentiated separately into preosteoblasts by day 14, with continued culture of these cells producing a mineralised osteogenic matrix, with a modified protocol able to produce cardiomyocytes. Though osteogenic differentiation of hPSC has been achieved by many different groups [48–50], we used this as proof of principal that the limb-bud progenitors are able to generate bone as well as an articular cartilage lineage (no COLX but PRG4, ACAN and COL2A1 expressing cells). In the long term this may allow development of osteochondral grafts produced from a unified differentiation process. This requires formation of a true osteochondral system complete with a mineralised cartilage boundary, which will ultimately allow for better integration of the graft and its long-term survival in situ. This would reduce the complexity of using multiple cell populations. Finally, whilst the RAPID protocol is designed to generate OCP and subsequent prechondrocytes and preosteoblasts, we showed the capability to generate beating cardiomyocytes. Confirming the lateral plate lineage by progressing to through the cardiac mesoderm [27], and including BMP as used by Takei *et al* [51] to deriving beating cells.

In conclusion, we have demonstrated the fast and efficient production of limb-bud/ articular progenitors which are able to form both cartilaginous and osteoid tissues with appropriate culture using a defined growth factor and small molecule regimen. The production of both cell types from the lateral plate lineage could be important in aiding graft incorporation by facilitating the formation of both the chondral and subchondral tissue, and generating the appropriate matrix required for hyaline cartilage. It is now important, as with paraxial differentiation methods [15], to understand the progress of transcriptome changes which occur during this protocol as identified in large RNAseq studies and the likely subtly different cell types generated providing matrix distribution heterogeneity. Comparison of these methods may indicate additional important factors associated with joint cartilage development.

## Acknowledgements

This work was supported by Arthritis Research UK (Grants R20786) and an MRC UKRMP hub award (Grant MR/K026666) to SJK as well as a Rosetrees Trust grant A1984 to CAS, SJK and Dr Mona Dvir-Ginzberg and studentships from the Engineering and Physical Sciences Research Council (EPSRC) and Medical Research Council (MRC): Centre for Doctoral Training (CDT) in Regenerative Medicine (EP/L014904/1) to PH, MN and FM. We thank Mr. Peter Walker for histology training. Aggrecan G1 antibody was generously donated by Prof Tim Hardingham of University of Manchester.

