## Supplementary data for "Directed differentiation of hPSCs through lateral plate mesoderm for generation of articular cartilage progenitors"

**Media compositions**

| **DDBM** | **Directed differentiation basal media** | |
| --- | --- | --- |
| **Component** | **Stock Conc** | **DDBM (final conc)** |
| DMEM/F12 (Sigma, D6421) | NA | NA |
| L-Glutamine (Gibco, 25030149) | 200mM | 5mL (2mM) |
| ITS (Gibco, 41400045) | X100 | 5mL (x1) |
| B27 (Gibco, A1895601) | X50 | 10ml (x1) |
| NEAA (Gibco, 11140050) | X100 | 5mL (x1) |
| Β-mercaptoethanol (Gibco, 31350010) | 50mM | 917uL (0.1M) |
| **CB** | **Chondrobasal media** | |
| **Component** | **Stock Conc** | **Chondro basal (final conc)** |
| DMEM (Sigma, D5796) | NA | NA |
| L-Glutamine | 200mM | 5mL (2mM) |
| Dexamethasone (Sigma, | 100uM | 500uL (100nM) |
| ASC-2-P (Sigma, A8960) | 5mg/mL | 5mL (50ug/mL) |
| Proline (Sigma, P5607) | 4mg/mL | 5mL (40ug/mL) |
| ITS | X100 | 5mL (x1) |
| **OB** | **Osteobasal media** | |
| **Component** | **Stock Conc** | **Osteo-basal (Final Conc)** |
| DMEM/F12 | NA | NA |
| L-Glutamine | 200mM | 5mL (2mM) |
| Dexamethasone | 100uM | 500uL (100nM) |
| ASC-2-P | 5mg/mL | 5mL (50ug/mL) |
| β-Glycerophosphate (Sigma G9422) | 1M | 10mM |
| ITS | X100 | 5mL (1x) |
| **BM** | **Basal-Cardio media** | |
| **Component** | **Stock Conc** | **Beta-A-Mix (Final Conc)** |
| DMEM/F12 | NA | NA |
| L-Glutamine | NA | 5mL (2nM) |
| Dexamethasone | 100uM | 5mL (100nM) |
| ASC-2-P | 5mg/mL | 5mL (50ng/mL) |
| B-Glycerophosphate | 1M | 10mM |
| ITS | X100 | 5mL (1x) |
| B27 | X50 | 10mL (1x) |
| NEAA | X100 | 5mL (1x) |
| β-mercaptoethanol |  | 917uL (0.1M) |

**List of QRT-PCR primer sequences**

| GENE | FORWARD | REVERSE |
| --- | --- | --- |
| ACAN | CATTCACCAGTGAGGACCTCG | TCACACTGCTCATAGCCTGCTTC |
| NKX2.5 | CAAGTGTGCGTCTGCCTTT | CAGCTCTTTCTTTTCGGCTCTA |
| ARID5B | GAGCAAAGGCATCTCCCAGT | GTTGAGGCCCGAGTTACACA |
| COL1A2 | CAGCCGCTTCACCTACAGC | TTTTGTATTCAATCACTGTCTTGCC |
| COL2A1 | GGCAATAGCAGGTTCACGTACA | CGATAACAGTCTTGCCCCACTT |
| TNNT2 | GGAGGAGTCCAAACCAAAGCC | TCAAAGTCCACTCTCTCTCCATC |
| GAPDH | ATGGGGAAGGTGAAGGTCG | TAAAAGCAGCCCTGGTGACC |
| GATA4 | CCTCCTCTGCCTGGTAATGACT | CGCTTCCCCTAACCAGATTG |
| GATA6 | GTGCCCAGACCACTTGCTAT | TGGGGGAAGTATTTTTGCTG |
| SPARC | ATCTTCCCTGTACACTGGCAGTTC | CTCGGTGTGGGAGAGGTAC |
| BGLAP | CTGGAGAGGAGCAGAACTGG | GGCAGCGAGGTAGTGAAGAG |
| NANOG | GGCTCTGTTTTGCTATATCCCCTAA | CATTACGATGCAGCAAATACAAGA |
| Brachyury | TGCTTCCCTGAGACCCAGTT | GATCACTTCTTTCCTTTGCATCAA G |
| MIXL1 | GGTACCCCGACATCCACTTG | TAATCTCCGGCCTAGCCAAA |
| CDX2 | GGGCTCTCTGAGAGGCAGGT | CCTTTGCTCTGCGGTTCTG |
| TBX6 | AAGTACCAACCCCGCATACA | TAGGCTGTCACGGAGATGAA |
| FOXF1 | AGCAGCCGTATCTGCACCAGAA | CTCCTTTCGGTCACACATGCTG |
| ISL1 | AGATTATATCAGGTTGTACGGGATCA | ACACAGCGGAAACACTCGAT |
| PDGFRa | CACACCTCCTCGCTGTAGTATTT | GTTATCGGTGTAAATGTCATCCA |
| PAX1 | CTCACAGCTGGCAGGGTATC | TTAGAGACTGCATGTTAGTTCTGGA |
| SOX5 | ATCCCAACTACCATGGCAGCT | TGCAGTTGGAGTGGGCCTA |
| SOX9 | GACTTCCGCCACGTGGAC | GTTGGGCGGCAGGTACTG |
| MEOX1 | TCAAAAGGAAGGAAATGACAGAGA | CCTCAGATGTGCAGCCTACAGA |
| SOX6 | GCAGTGATCAACATGTGGCCT | CGCTGTCCCAGTCAGCATCT |
| RUNX2 | GACGAGGCAAGAGTTTCACC | GCCTGGGGTCTGTAATCTGA |
| COLX | CACCAGGCATTCCAGGATTCC | AGGTTTGTTGGTCTGATAGCTC |
| PRG4 | GAGTACCCAATCAAGGCATTATCA | CCATCTACTGGCTTACCATTGCA |
| PRRX1 | TGATGCTTTTGTGCGAGAAGA | AGGGAAGCGTTTTTATTGGCT |
| HOXB5 | AACTCCTTCTCGGGGCGTTAT | CATCCCATTGTAATTGTAGCCGT |
| HAND1 | GTGCGTCCTTTAATCCTCTTC | GTGAGAGCAAGCGGAAAAG |
| HAND2 | ACATCGCCTACCTCATGGAC | TTCTTGTCGTTGCTGCTCAC |

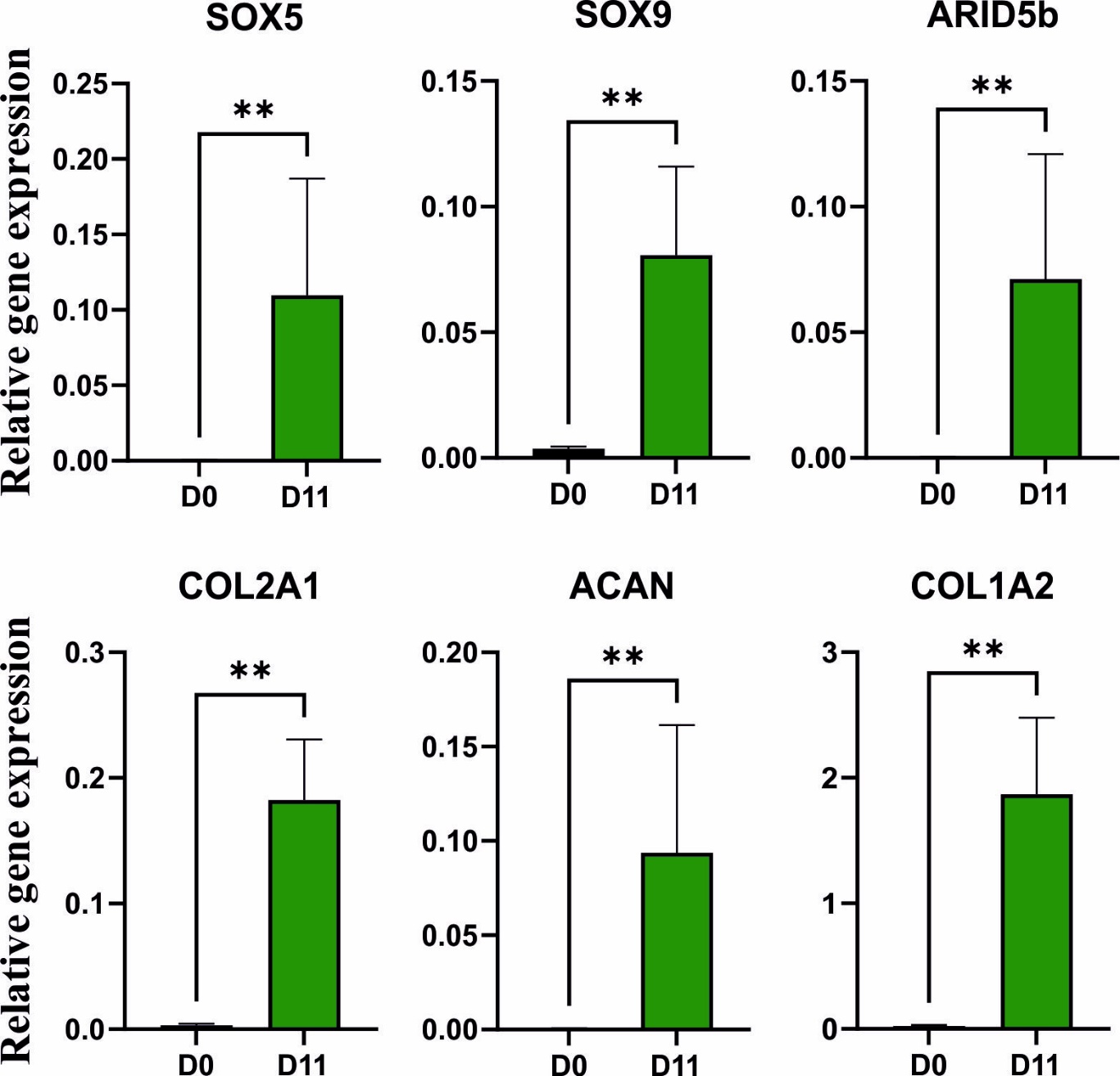

**Figure S1.** **Differentiation of Man7 hPSCs to chondroprogenitors**. Human embryonic stem cell line Man7 was differentiated through the RAPID protocol to produce chondroprogenitors with samples taken at Day 11 (D11). Gene expression was assessed by qRT-PCR for chondrogenic genes. Mann-Whitney test was used to determine statistical significance. Significance relative to hPSC (day 0) control cells (+ ≤0.05, ++≤0.01) (N=6 biological repeat).
